# CellKb Immune: a manually curated database of mammalian hematopoietic marker gene sets for rapid cell type identification

**DOI:** 10.1101/2020.12.01.389890

**Authors:** Ajay Patil, Ashwini Patil

**Affiliations:** Combinatics, Ichikawa-shi, Chiba, Japan

**Keywords:** single-cell, cell types, marker genes, immune cells, annotation, database, cell type prediction

## Abstract

Using single-cell RNA sequencing, it is now possible to get the detailed gene expression pattern in each cell for millions of cells in a single experiment under varying conditions. Annotating cells by assigning a type or state to them is important for correct interpretation of results. In spite of current advances in technology, assigning cell types in single-cell datasets remains a bottleneck due to the lack of a comprehensive reference database and an accurate search method in a single tool. CellKb Immune is a knowledgebase of manually collected, curated and annotated marker gene sets representing cell types in the immune response in mouse. It allows users to find matching cell types in literature given a list of genes. We evaluated the contents and search methodology of CellKb Immune using a leave-one-out approach. We further used CellKb Immune to annotate previously defined marker gene sets from a bulk RNA-seq dataset to confirm its accuracy and coverage. CellKb Immune provides an easy-to-use online tool with a reliable method to find matching cell types and annotate cells in single-cell experiments. It is available at https://www.cellkb.com/immune.

## 1 Introduction

Single-cell RNA-seq is widely used to study transcriptional patterns of genes in individual cells. Various computational tools have been developed for the annotation of cell types to cells in single-cell experiments, given a reference database and a gene list (1,2). Several single cell reference datasets or atlases have also been prepared (3–5), or compiled into aggregate reference databases of cell types and marker gene sets (6–9). There are currently two primary methods used for annotating cells in single-cell experiments. 1) Manually searching through scientific literature to find marker genes associated with cell types being studied. This requires prior knowledge of the cell types potentially present in the sample and fails to identify rare cell types without established markers. 2) Selecting a single cell reference database/ dataset from literature, downloading and processing the data, identifying and installing a computational method to assign cell types, and finally, writing a program in R or Python to perform the cell type assignment. This method usually limits a researcher to using a single, or a few datasets as a reference. Both these methods require significant time and effort but fail to use the vast amount of cell type marker gene sets previously published. Reliable annotations require a resource that contains a comprehensive reference database of cell types and marker gene sets from literature along with a search method that allows users to find matching cell types.

CellKb Immune addresses these issues by collecting and aggregating a large amount of immune-related cell type signatures in mouse from literature and providing an online interface to find matching annotated cell types easily, quickly and accurately.

## 2 Materials and Methods

### 2.1 Data collection and curation

The CellKb Immune database consists of marker gene sets of hematopoietic cell types with associated annotations collected from literature. The marker genes and their cell type annotations are taken primarily from published single-cell RNA-seq experiments. All publications related to single-cell experiments are taken from PubMed and manually screened to select those identifying cell type specific gene expression patterns.

Cell type specific marker gene sets are defined as those identified by authors based on their significant change in expression within a group of cells as compared to all other cells in the experiment. The marker gene sets are extracted from supplementary materials and raw, or processed, data of publications describing these experiments. If calculated based on the raw/processed expression data, cell type specific marker genes are identified using the methods described by the authors in the publication, along with cell type annotations given for the single cells.

Marker gene sets are selected from published experimental studies if they satisfy the following criteria: 1) Raw data related to the experiment is deposited in public databases, 2) The marker gene sets are publicly available for download, 3) The experimental methods and conditions are adequately described in the manuscript to allow detailed annotation, 4) The number of cells and samples studied is clearly specified, 5) The computational methods used to normalize, filter and cluster cell types, along with identification of cluster-specific genes is clearly described, 6) Associated values such as average expression, fold change, statistical significance, raw UMI or read counts, are available for marker genes (marker gene sets specified only as ranked gene lists without associated values are included in some cases), 7) A majority of the gene identifiers in the marker gene set are valid and can be mapped to the latest version of the Ensembl database (10). Single-cell experiments without cell type annotations are not used to calculate cell type specific marker gene sets.

Detailed sample and treatment conditions, including sample counts and cell counts are also stored in CellKb Immune. Information about disease state of the samples and the cell types enriched in disease samples are also noted. All data in CellKb Immune is referenced and linked to the PubMed identifier of the source publication.

CellKb Immune also contains marker genes extracted from select bulk RNA-seq and microarray experiments from public databases. These include selected gene signatures from MSig-db (11) and previously published bulk RNA-seq datasets of immune cell types (1,12). Cell-type specific interaction networks are created using marker gene sets of all cell types and protein-protein interactions with reliability scores from the HitPredict database (13).

### 2.2 Data harmonization and annotation

Data harmonization is essential for rapid search and retrieval of matching cell types. Given that there is no standard format currently in place for sharing cell type specific marker gene sets, CellKb Immune performs a number of operations to harmonize the data collected. Marker gene sets are extensively curated to identify valid genes and cell types. Genes given in the published marker gene sets are mapped to valid Ensembl identifiers by matching names, synonyms or identifiers. Assigning a common gene identifier to the marker genes is an important part of data harmonization in CellKb Immune.

Associated values given by authors with each signature are also stored in CellKb Immune. These include, but are not limited to, rank, score, average expression, log fold change and corrected/uncorrected p-values.

For each marker gene set up to 1,500 marker genes with corrected p-values less than or equal to 0.1 and log fold change greater than 0 compared to other cell types are selected. The marker genes are ranked by CellKb Immune based on descending log fold change values. Marker gene sets that are published in the form of ranked gene lists without associated values are also included in the database if they match an existing marker gene set with the same cell type annotation. Unranked gene lists without associated values are not included in CellKb Immune.

Each marker gene set is associated with a cell type, tissue/organ and disease condition given in the source publication, which are mapped to standardized ontology terms using Cell Ontology (14), Uberon Ontology (15) and Disease Ontology (16).

Each marker gene set is assigned a reliability score that gives an indication of how similar it is to other marker gene sets of the same ontology. The higher the reliability score of a marker gene set, the closer it is to other marker gene sets of the same cell type and, hence, the more likely it is to be accurate. This helps identify marker gene sets that are incorrect, calculated from erroneous cell cluster assignments by authors, or that have been calculated from mixtures of cell types.

Finally, all marker gene sets and their annotations are stored in a uniform format with indexed identifiers enabling rapid search and retrieval of the data.

### 2.3 Data searching

In CellKb Immune, users can search for cell type specific marker gene sets by giving a list of statistically significant query genes ranked in descending order of log fold change. Query genes given by the user are compared with every marker gene set in the database and the best match is identified using a rank-based match score.

The match score is based on the number of common genes between the query and a marker gene set, their ranks, their rank differences and the total number of significant genes in the marker gene set. This results in marker gene sets sharing highly ranked genes with the query being assigned higher match scores. This method also accounts for differences in the sizes of gene lists between the query and various marker gene sets, such those with fewer marker genes are not disregarded.

## 3 Results

### 3.1 Database contents

CellKb Immune contains author-defined mouse hematopoietic cell type marker gene sets manually collected from publications describing mainly single-cell, and selected bulk RNA-seq or microarray experiments (1,11) (Figure 1). Cell type marker gene sets are assigned a Cell Ontology term (14), where available. Tissues, organs or body parts of the experimental sample are assigned Uberon ontology terms (15). Disease ontology terms are provided where the marker gene sets are obtained from a diseased sample (16). All marker gene sets reference their source publication with a link to its PubMed identifier.

**Figure 1.**
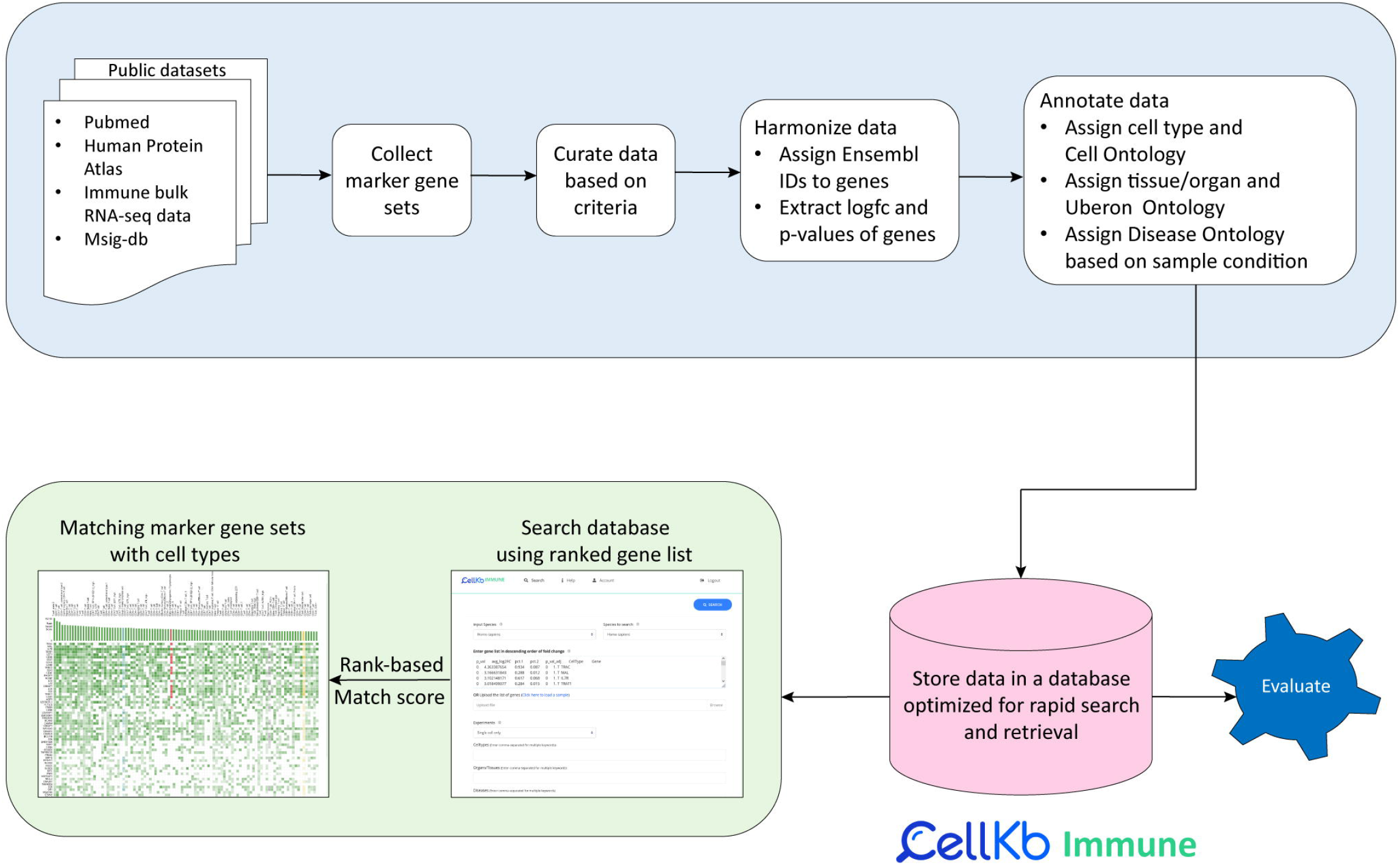
CellKb Immune workflow – data collection, harmonization and search interface.

CellKb Immune contains:

- 2,395 marker gene sets from mouse
- 230 unique annotated hematopoietic cell types (Cell Ontology CL:0000988 and all its descendants).
- 104 tissues/organs
- 226 publications selected from approximately 10,000 studies published between 2013 and 2021.
- 40,122 protein-protein interactions from HitPredict (13)

### 3.2 Functionality

The CellKb Immune web interface allows users to search matching cell types and their marker gene sets in mouse. It can be used to annotate cells from single-cell RNA-seq data, given a list of statistically significant cluster-specific genes ranked by decreasing log fold change. Associated values such as log fold changes and p-values are also accepted as input by CellKb Immune. Genes in the user query are compared with every marker gene set in the database using a rank-based search method and a match score is computed (see Methods). The user is notified if the query gene list matches non-hematopoietic cell types in 5 out of top 10 marker gene set matches. Figure 2 shows the CellKb Immune user interface.

**Figure 2.**
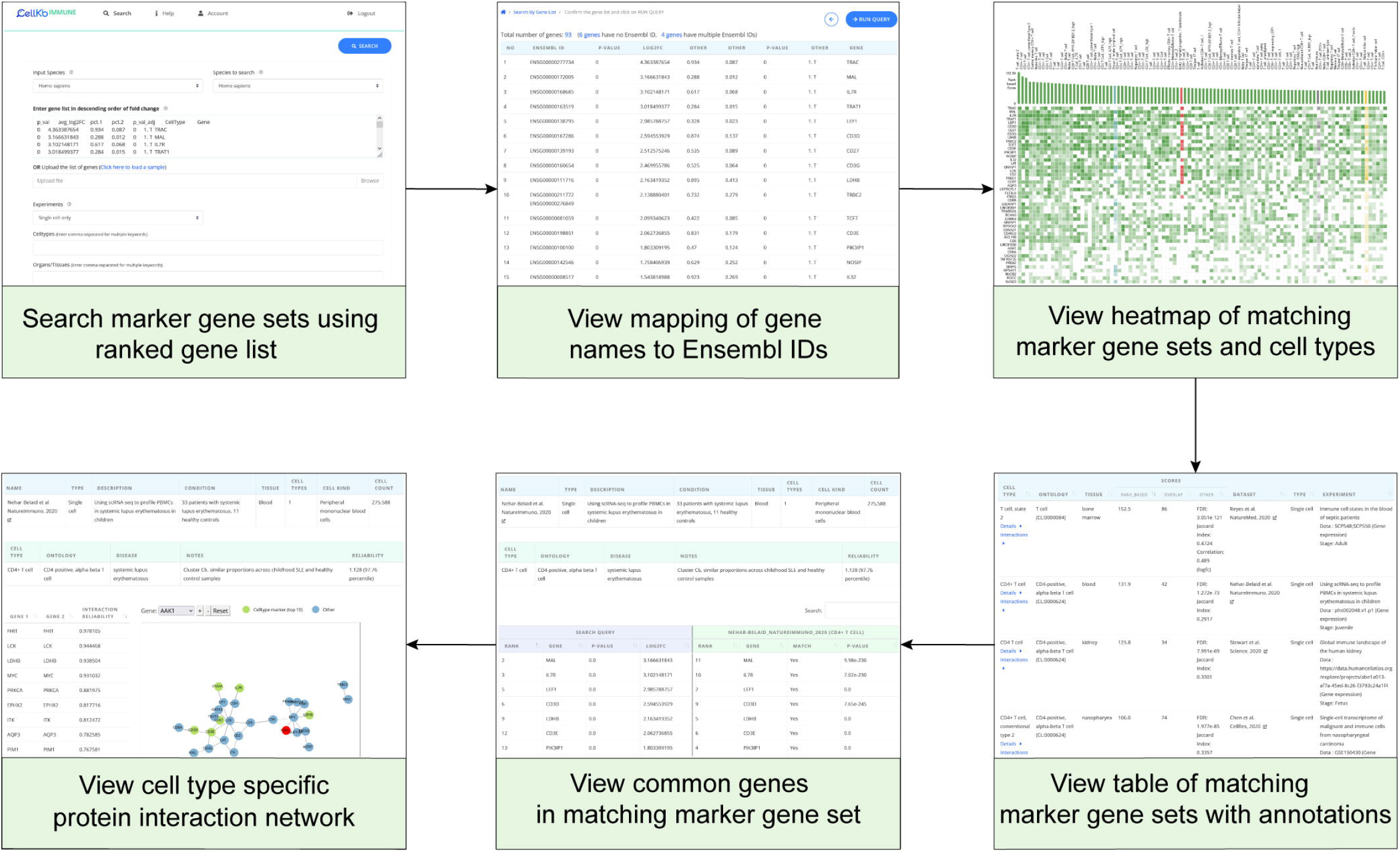
CellKb Immune user interface used to search matching cell types given a gene list.

### 3.3 Evaluation

To check the database quality and cell type matching performance of the CellKb Immune search method, we used the leave-one-out approach. We matched each of the 2,223 mouse marker gene sets annotated with cell ontologies in CellKb Immune with all others in the database. 1,568 (70.5%) of gene sets were correctly matched to another gene set with the exact same ontology or its parent, while for 427 (19.2%) gene sets, the ontology of the first hit was a sibling i.e. a descendant of the same parent ontology (Table 1). Thus, 89.7% of marker gene sets were matched to another marker gene set with the same ontology, or its parent or sibling (Table S1).

**Table 1.** Performance evaluation of CellKb Immune

| Evaluation database | Marker gene sets | CellKb Immune performance |  | Total ontologies matched |
| --- | --- | --- | --- | --- |
|  |  | Correct ontology match | Parent ontology match |  |
| CellKb Immune Mouse (leave-one-out) | 2,223 | 1,568 (70.5%) | 427 (19.2%) | **1,995 (89.7%)** |
| Yoshida et al. (12) Mouse (all-against-all) | 98 | 61 (62.2%) | 18 (18.4%) | **79 (80.6%)** |

We further evaluated the performance of the CellKb Immune by checking the number of correct cell types assigned to gene lists obtained from a bulk RNA-seq dataset of mouse immune cell types (12) from Immgen (17). Using the top1,500 significantly upregulated genes for each cell type in the bulk RNA-seq dataset, we searched the CellKb Immune database for matching marker gene sets. We then compared the cell ontology of the top 10 hits with that of the ontology assigned to the gene lists in the query dataset. Of the 98 marker gene sets representing 40 distinct cell types, CellKb Immune was able to find the exact matching ontology or its parent as the first hit in 61 instances (62.2%). In another 18 marker gene sets (18.4%), the ontology of the first hit was a sibling i.e. a descendant of the same parent ontology as that in the query dataset (Table 1). Thus, a total of 79.5% of marker gene sets in the bulk RNA-seq dataset had their first hit as the same ontology, its parent or sibling ontology (Table S2).

These results show that 1) the CellKb Immune database includes high quality annotated marker gene sets from single-cell experiments, and 2) the search methodology used by CellKb Immune to find matching cell types is able to recapitulate the original cell types from within its own database and other unseen bulk and single-cell datasets.

We next compared the functionality of CellKb Immune to two other similar databases, CellMarker (7) and PanglaoDB (8) (Table 2). CellKb Immune is unique in its ability to search matching cell types from published literature using ranked gene lists. CellMarker allows users to search only one gene at a time, while PanglaoDB can search cell types using unranked gene lists. Additionally, both CellMarker and PanglaoDB contain gene signatures that have been aggregated from multiple samples and/or sources. In contrast, CellKb Immune contains cell type marker gene sets as described in literature and gives the overlapping genes between the query and the matching cell types obtained in a search result. Some other features that differentiate CellKb Immune from CellMarker and PanglaoDB are deep annotations in the form of standardized ontologies and protein-protein interaction networks. CellKb Immune also contains marker gene sets recently published in literature and is regularly updated.

**Table 2.** Comparison of CellKb Immune functionality with other similar tools.

| **Feature** | **CellKb Immune** | **CellMarker (7)** | **PanglaoDB (8)** |
| --- | --- | --- | --- |
| Last update | 10 February 2022 | 3 November 2018 | 27 March 2020 |
| Manual curation | Yes | Yes | No |
| Ranked marker genes with associated values, eg. log fold change, FDR | Yes | No | Yes |
| Marker gene sets per cell type | Multiple (specified as given in literature) | Single (aggregated from multiple sources) | Single (aggregated from multiple sources) |
| Marker genes obtained from single-cell RNA-seq experiments | Yes | Yes | Yes |
| Processing of single-cell RNA-seq data to calculate marker genes | Using author-specified data processing pipeline | No | Using standardized data processing pipeline |
| Marker genes obtained from bulk RNA-seq/ microarray experiments | Yes | Yes | No |
| Disease information | All | Cancer only | All |
| Protein-protein interaction networks of marker genes | Yes | No | No |
| Search matching cell types with ranked gene lists | Yes | No | No |
| Cell/Uberon/Disease ontology annotations | Yes | Limited Cell Ontology annotations | No |
| Reliability score for marker gene sets | Yes | No | Yes |

Thus, CellKb Immune provides complementary contents and functionality to other cell type marker databases.

## 4 Discussion

CellKb Immune has been designed as a resource to find matching cell type specific marker gene sets from literature to help assign cell types to cells in single-cell RNA-seq experiments. CellKb Immune addresses several issues in existing single-cell reference databases and tools used for this purpose.

CellKb Immune consists of user-defined marker gene sets from literature. These carry significant knowledge since the cell types in the form of cell clusters are often selected by authors based on biological information. CellKb Immune is able to capture and aggregate this biological information. On the other hand, several reference databases reanalyze the raw expression patterns using a common pipeline while ignoring the methodology and cluster definitions used by the authors in the original study, thus losing valuable biological information.

CellKb Immune provides deep annotations, in the form of extensive cell type information and description, curated and validated marker genes ranked by significance, along with fold change and significance values associated with gene expression. Additionally, CellKb Immune provides a web-based interface to find matching cell types in published data sets given a user gene list, independent of the experimental platform, analysis methods and size of the marker gene sets. Thus, users do not need to spend time integrating reference data and search methods programmatically.

Unlike other computational methods which require the presence of associated expression or fold change values and the same number of genes in all target cell types, the rank-based search method used by CellKb Immune enables searching through marker gene sets of differing sizes, in the absence of expression fold changes, and independent of experimental platform and pre-processing methods. Searching of cell types happens by direct comparison of the query gene list with the marker genes of every cell type in CellKb Immune using a fast and reliable method without any type of score aggregation or machine learning. As a result, CellKb Immune provides information about the matching cell type in the form of the publication, experimental details and the overlapping genes between the query and matching marker gene set. This allows users to further explore the study to which their cell type matches. In cases where the top matches to a marker gene set from single-cell data are inconclusive, users can refer to the consensus of the top matching cell types or compare their gene lists with gene sets from bulk RNA-seq or microarray experiments.

Since CellKb Immune contains marker gene sets from multiple publications for each cell type, it can also be used to evaluate the validity of known marker genes for the cell type of interest. The reliability score assigned to each cell type marker gene set can also be used to identify those that are incorrectly annotated in publications. Additionally, comparing the query gene list with marker gene sets from multiple publications helps assess the accuracy of the matching cell types found by CellKb Immune.

Thus, CellKb Immune provides an easy-to-use reference database with a fast and reliable method to find matching cell types and marker gene sets to help annotate cells in single-cell experiments in a single tool.

## 6 Author Contributions

Ashwini conceived of the project, collected the data and prepared the manuscript. Ajay and Ashwini designed and developed the database, search method and web application. All authors read and approved the final manuscript.

## 10 Supplementary Material

Table S1. Results of the leave-one-out evaluation in CellKb Immune shows the first hit of each cell type marker gene set compared to all other marker gene sets within the species.

Table S2. First hit of each cell type marker gene set in a bulk RNA-seq dataset of immune cell types compared to all marker gene sets in CellKb Immune.

## 11 Data Availability Statement

CellKb Immune is available at https://www.cellkb.com/immune. All data used in CellKb Immune is publicly available and appropriately referenced in the web application.

